# Comparison studies between Cesium-137 and X-ray irradiators in epithelial injury using *in vitro* and *in vivo* models

**DOI:** 10.64898/2026.04.17.719248

**Authors:** Rabina Lakha, Emilia J. Orzechowska-Licari, Sahaana Kesavan, Zhi J. Wu, Matthew Rotoli, Michael Giarrizzo, Vincent W. Yang, Agnieszka B. Bialkowska

## Abstract

Radiation-induced intestinal injury is a widely used model for studying mechanisms regulating tissue injury and regeneration. Traditionally, Cesium (^137^Cs) radiation has been used in research applications, but over the past decade, X-ray irradiation has become increasingly favored due to its improved safety and non-radioactive profile. Since each type of radiation has distinct physical characteristics that drive its performance, we sought to systematically compare the effects of the X-ray and ^137^Cs irradiators on intestinal epithelial injury and regeneration. Using established *in vitro* models, including colorectal cancer cell lines such as HCT116, RKO, and DLD-1, and mouse intestinal organoids, alongside *an in vivo* model, *Bmi1-CreER*;*Rosa26eYFP*, we evaluated differences in transcriptional, protein, and histopathological responses to irradiation. Our results demonstrate that X-ray produced intestinal injury and regenerative responses comparable to those induced by ^137^Cs, supporting its reliability as an alternative modality for studying intestinal radiation.

## Introduction

Studies concerning post-irradiation intestinal injury in mouse models have traditionally been performed using gamma-emitting cesium-137 (^137^Cs) irradiators^1^. However, following the establishment of the Cesium Irradiator Replacement Project (CIRP) by the Department of Energy (DOE) in 2014 due to national security risks associated with ^137^Cs irradiators, a significant global effort has been underway to replace ^137^Cs irradiators with safer, non-radioactive X-ray alternatives^2^. ^137^Cs irradiators undergo continuous radioactive decay to produce gamma rays, are very expensive, and require a strict protocol for use and deployment. At the same time, X-ray irradiators are mostly safe, low-maintenance, electronically controlled, and easy to dispose of^2^.

While clear security and safety reasons support a move toward an X-ray irradiator, from a research perspective, there are important physical properties of these two irradiation sources that may affect the outcomes of experiments^3,4^. It has been shown that X-rays can have a stronger biological effect than ^137^Cs-irradiators. In contrast, ^137^Cs irradiation provides deeper penetration and more uniform dosing for larger samples, which can account for differences in tissue injury^5^. Comparison studies of bone marrow in different mouse models using ^137^Cs and X-ray irradiators have shown similar effects, but with important differences in physiological responses^4,6–8^. All these studies conclude that more defined, comprehensive studies with individual optimization in each field are necessary largely due to differences in the radiation sources between used with ^137^Cs and X-ray irradiators^9^.

Very few studies have been performed to characterize intestinal injury and regeneration after X-ray irradiation, and/or performed preliminary comparative studies with ^137^Cs irradiators^10^. As such, it is of utmost importance to assess the reproducibility of tissue injury and the mechanisms of repair induced by X-ray irradiation.

The intestinal epithelium is one of the most studied models for homeostasis and regeneration after radiation injury, due to its self-renewal capacity^11,12^. The entire epithelium is replaced in approximately 3-5 days as a consequence of the presence of active intestinal stem cells (aISCs) and reserve intestinal stem cells (rISCs), and their progenitors^13^. aISCs are present at the base of the crypts, maintaining homeostasis in the intestinal epithelium, and are generally marked by LGR5^14^, ASCL2^15^, OLFM4^16^, PROM1^17^, and SOX9^low18^. rISCs replenish aISCs following ionizing radiation to restore tissue architecture and functions and are marked by BMI1^19,20^, mTERT^21^, HOPX^22,23^, LRIG1^24,25^, SOX9^high18^, and KRT19^26^. Following irradiation, the regenerative response of the small intestinal epithelium has been categorized into three distinct phases: apoptotic, regenerative, and normalization. The apoptotic phase ranges from 24 to 48h post-irradiation. It is characterized by cell death specifically within the crypt regions and shortening of the villi. In contrast, the regenerative phase begins at 72 hours and culminates with the normalization at seven days post-injury^27–31^. Multiple studies investigating mechanisms of radiation injury responses, using either *in vitro* or *in vivo* models, have been conducted predominantly with ^137^Cs irradiators. Thus, in this study, we compared ^137^Cs and an X-ray irradiator to assess their effects on three well-established colorectal cancer cell lines, enteroids derived from mouse small intestine, and two mouse models. Our studies showed that on the macroscopic level, ^137^Cs and X-ray irradiators performed similarly.

## Results

### Colorectal cancer cell lines

Three colorectal cancer (CRC) cell lines (HCT116, RKO, and DLD-1) were irradiated with 12Gy using a ^137^Cs or X-ray irradiator (Supplementary Fig. 1a). Samples were collected at 6-, 24-, 48-, and 72-hour post-irradiation, and RNA and protein analyses were performed. Our data show that the levels of *CDKN1A* are readily induced upon irradiation in three tested cell lines (Figure 1a, f, and k), and these levels remain increased over a three-day experiment, regardless of the source of irradiation. The levels of *KLF5*, *KLF4*, *BAX*, and *BCL2* (Figure 1b-o) are altered upon radiation; however, they follow a similar pattern of modification (decrease or increase) within each CRC cell line, despite the source of irradiation.

**Figure 1.**
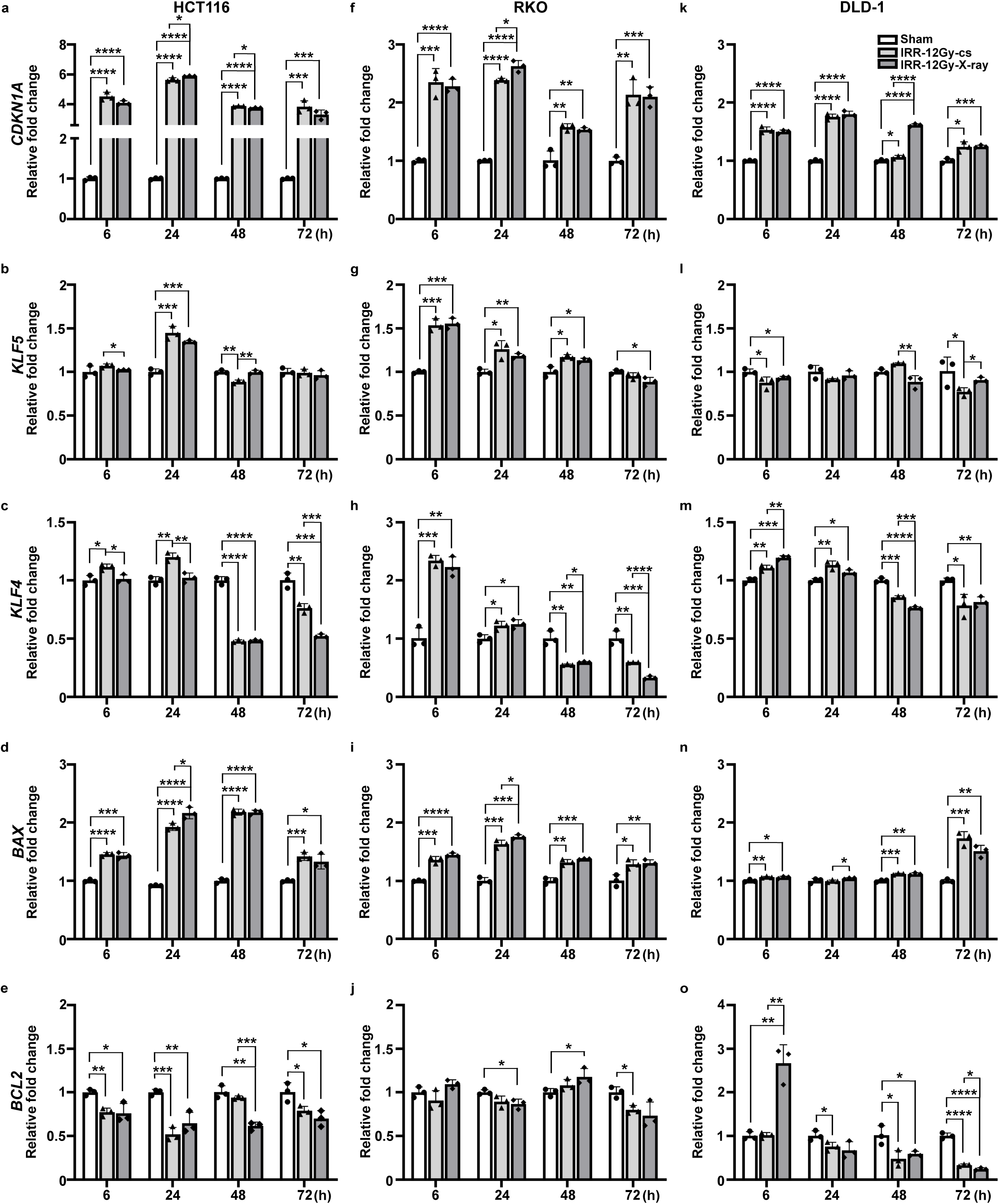
Response to injury in three colorectal cancer cell lines after 12Gy irradiation using ^137^Cs or X-ray irradiators. RT-qPCR analysis of *CDKN1A* (**a, f, k**), *KLF5* (**b, g, l**), *KLF4* (**c, h, m**), *BAX* (**d, i, n**), and *BCL2* (**e, j, o**) in HCT116, RKO and DLD-1 calculated as a fold change at 6, 24, 48 and 72 h after irradiation. *HPRT1* was used as a housekeeping gene (control). RT-qPCR was performed as described in the “Materials and methods” section. Data points represent the average of three independent experiments, with the mean ±SD indicated. Significance was determined by the Student’s test followed by an analysis of the normal distribution (Tukey’s test), *p < 0.05, **p < 0.01, ***p < 0.001, ****p < 0.0001.

For Western blot analysis, we assessed several markers of cell injury, including p21, γH2AX, TP53, and phospho-TP53 (Figure 2 and Supplementary Figure 2). Under both irradiation conditions, the levels of these markers significantly increased upon injury compared with sham-irradiated cells and persisted for at least 72 h in the HCT116 cell line. We observed increases in TP53, phospho-TP53, and p21, with a trend toward increased γH2AX in the RKO cell line under both radiation conditions. At the same time, DLD-1 cells responded with a significant increase in phospho-TP53, with a minimal change to P53 or p21. Additionally, we assessed the levels of pro-apoptotic marker BAX (BAX), as well as pro-proliferative marker - Krüppel-like factor 5 (KLF5) (Figure 2 and Supplementary Figure 2). We have not noticed differences in the response between the tested sources of irradiation.

**Figure 2.**
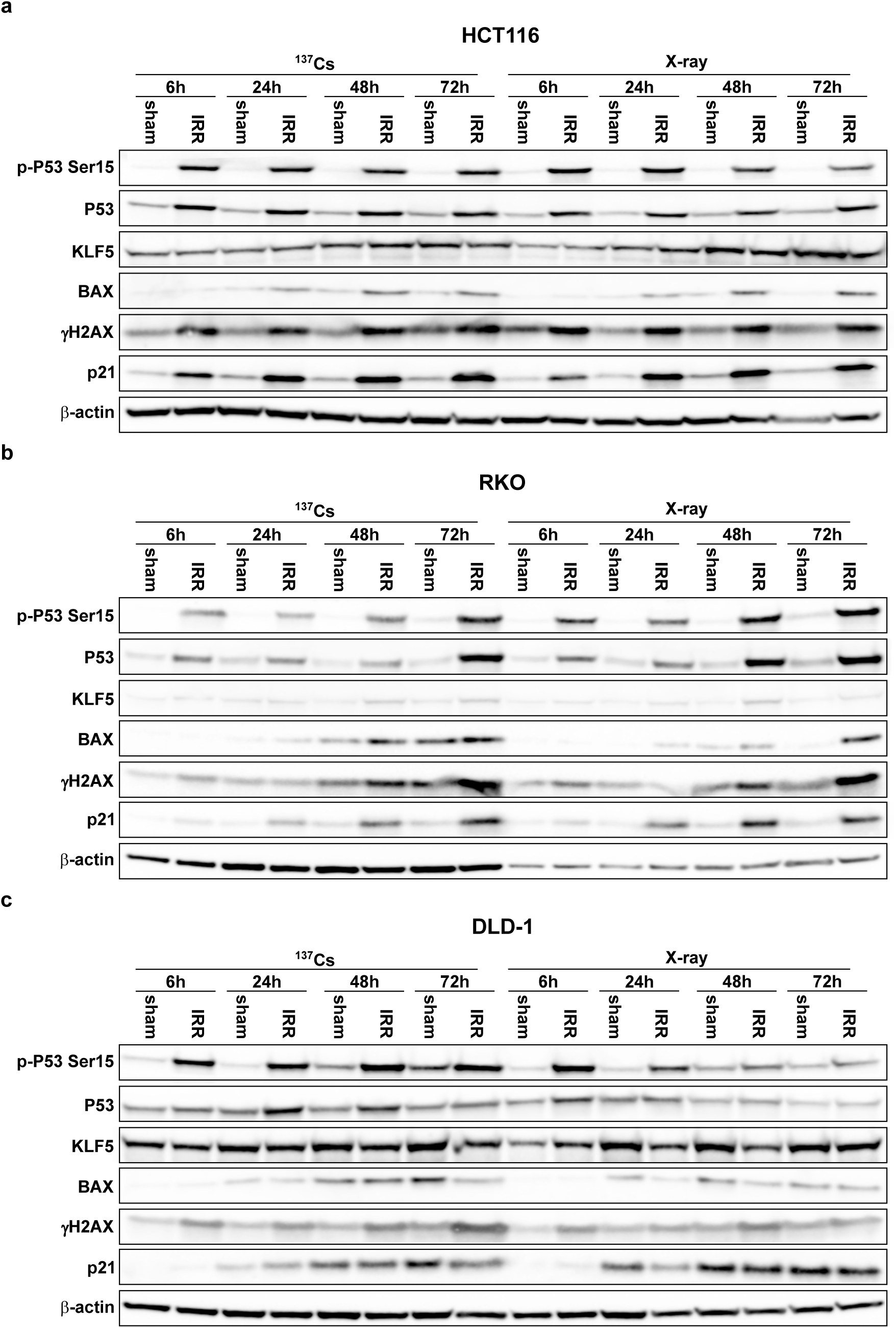
Western blot analysis of markers of cell injury, proliferation, and survival in three colorectal cancer cell lines exposed to 12Gy irradiation using ^137^Cs or X-ray irradiators. Markers of cell injury (p21, γH2AX, TP53, and phospho-TP53), proliferation (KLF5), and pro-apoptotic (BAX) markers were analyzed using Western blot for HCT116 (**a**), RKO (**b**), and DLD-1(**c**) at 6-, 24-, 48-, and 72 h post-irradiation. The figure displays representative images from *N = 3*.

### Mouse intestinal enteroids

Intestinal enteroids derived from *Bmi1-Cre^ER^* mice were cultured in L-WRN conditioned media and exposed to 6Gy or 8Gy irradiation using ^137^Cs or X-ray irradiators (Supplementary Figure 1b). The enteroids were imaged over 4 days after irradiation (Figure 3), and samples for downstream analysis were collected at 6, 24, 48, 72, and 96 h. Both 6Gy and 8Gy irradiation using either ^137^Cs or X-ray irradiator resulted in loss of budding in enteroids at 24 and 48h, indicating an apoptosis phase followed by new budding formation at 96h, related to the regeneration of organoids (Figure 3). For qRT-PCR analysis, we assessed the expression levels of *Cdkn1a* (injury marker), *MKi67* (proliferation marker), *Lgr5* and *Olfm4* (active stem cell markers), *Alpi* (absorptive intestinal epithelial cell marker), and *ChgA* (enteroendocrine intestinal epithelial cell marker) (Figure 4). *Cdkn1a levels* were increased at the tested time points regardless of the source or strength of irradiation and exhibited similar expression patterns (Figure 4a and b). However, we observed significant differences in the patterns of *MKi67*, *Lgr5*, and *Olfm4* expression levels. *MKi67* levels were significantly increased at 6h after 6Gy, regardless of the irradiation source, and steadily decreased at 24 and 48 hours. At 96 h post-irradiation, we observed a significant increase in *MKi67* transcript, but only in enteroids exposed to the X-ray irradiator. Similarly, 8Gy caused the same effect at 96 h on *MKi67* levels (Figure 4c and d). The strength of the irradiation did not affect the pattern of the expression of *Lgr5* at 6, 24, and 48h post-injury. It showed significant differences at 96 hours, with *Lgr5* transcript levels increasing after X-ray irradiation compared to ^137^Cs (Figure 4e and f). Similarly, *Olfm4* levels were significantly higher after irradiation of enteroids with the X-ray irradiator than with the ^137^Cs irradiator (Figure 4g and h). In addition, we observed differences at the early time point (6h) in the levels of *Mki67*, *Lgr5*, and *Olfm4*; however, these differences were due to the strength of the irradiation (8Gy versus 6Gy), not its source. We noticed that absorptive cell marker (*Alpi1*) levels decreased when enteroids were irradiated with ^137^Cs, whereas they were unaffected or slightly increased with the X-ray irradiator (Figure 4i-j). In contrast, we observed that the changes to the levels of the tested enteroendocrine cell maker (*Chga*) did not depend on the strength of the irradiation (6Gy versus 8Gy) (Figure 4k-l).

**Figure 3.**
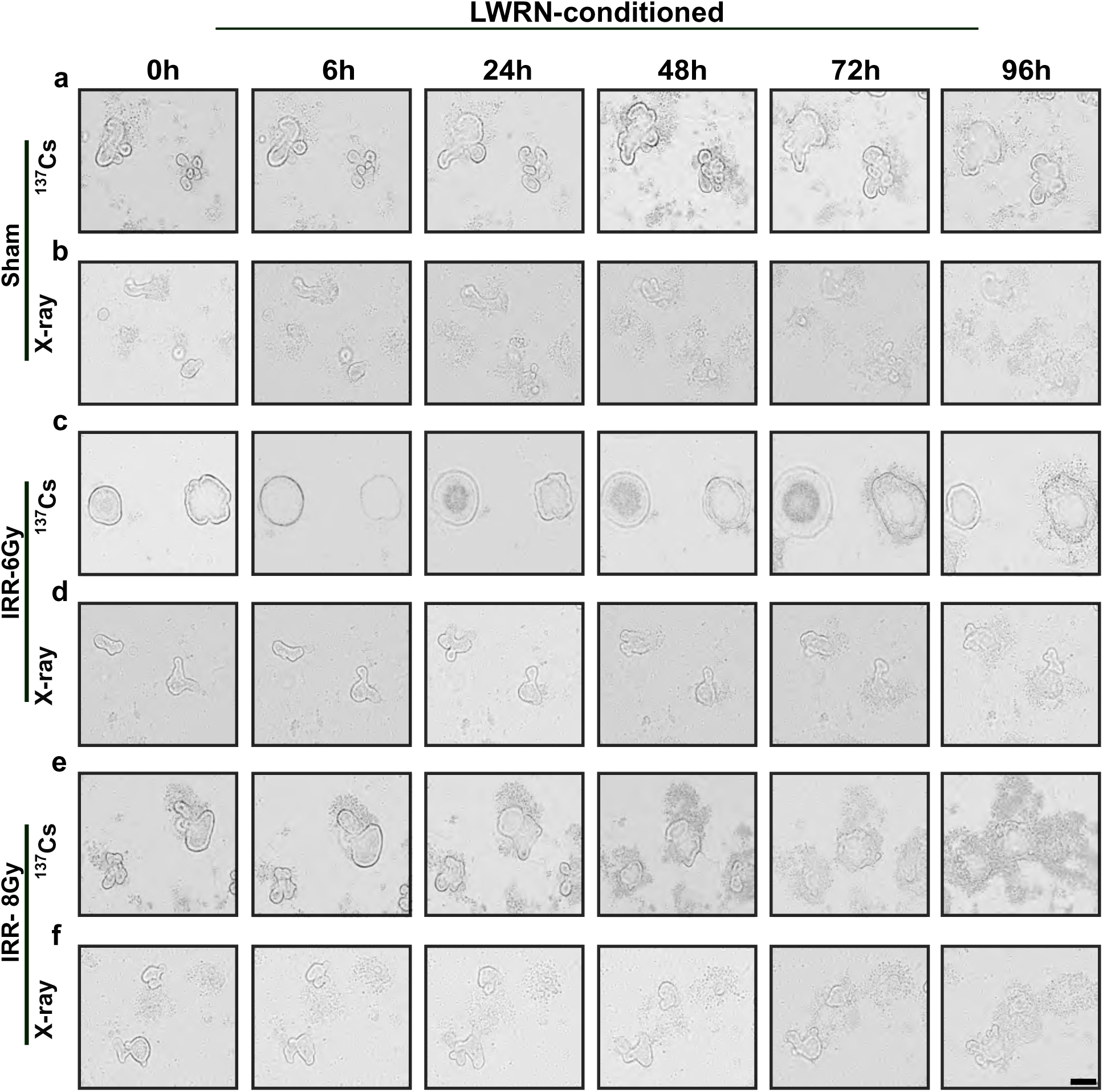
Brightfield images of *Bmi1-Cre^ER^*-derived organoid after 6Gy or 8Gy ^137^Cs or X-ray irradiation. Representative time-course images were obtained from sham-irradiated (**a-b**), 6Gy-irradiated (**c-d**), or 8Gy-irradiated (**e-f**) at 0-, 6-, 24-, 48-, 72-, and 96 h post-irradiation. The figure displays representative images from *N = 3*. Scale bar = 200 µm.

**Figure 4.**
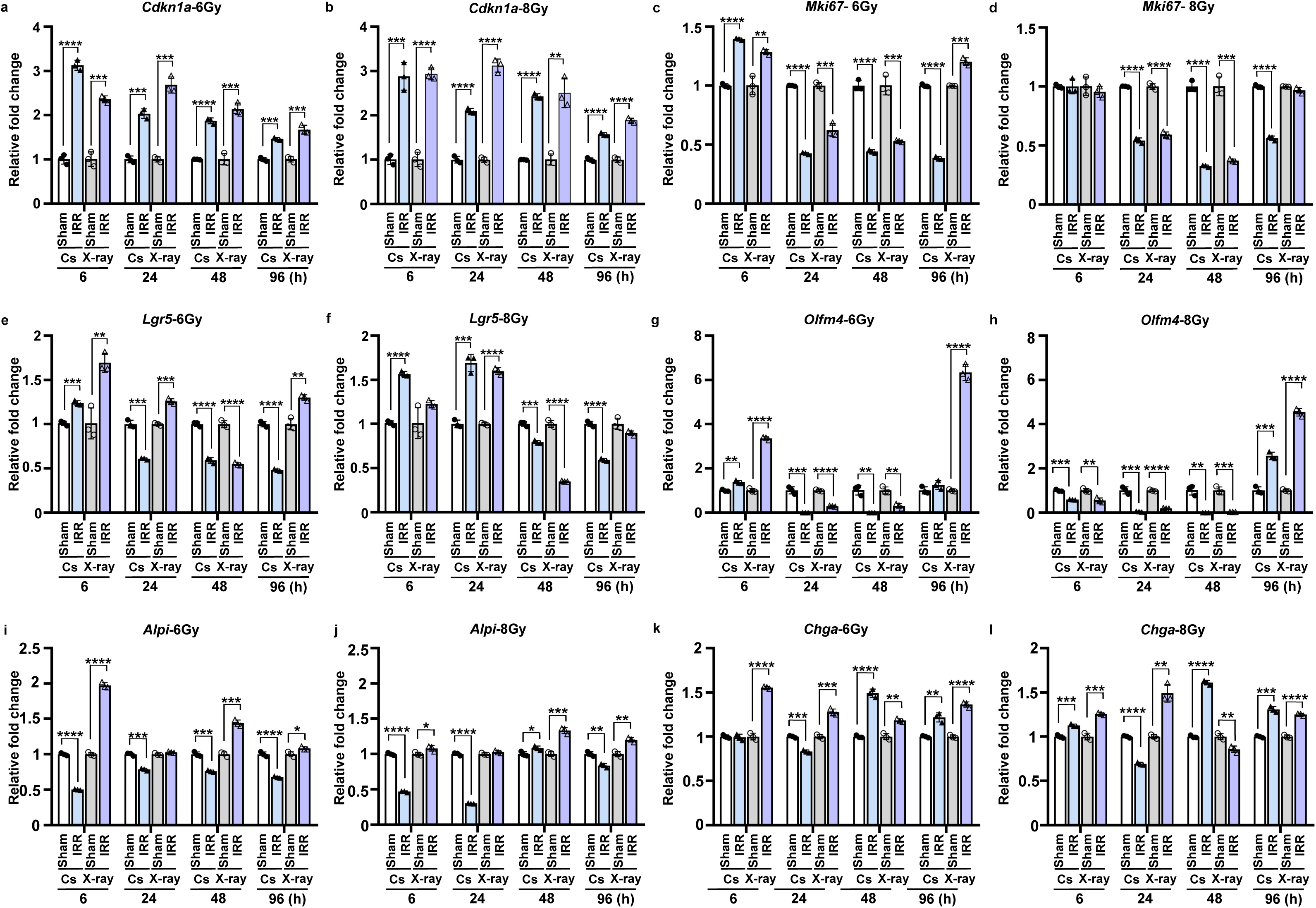
RT-qPCR analysis of *Bmi1-Cre^ER^*-derived enteroids after 6Gy or 8Gy ^137^Cs or X-ray or sham irradiation. RT-qPCR analysis of (**a**) *Cdkn1a* expression levels after 6Gy, (**b**) *Cdkn1a* expression levels after 8Gy, (**c**) *Mki67* expression levels after 6Gy, (**d**) *Mki67* expression levels after 8Gy, (**e**) *Lgr5* expression levels after 6Gy, (**f**) *Lgr5* expression levels after 8Gy, (**g**) *Olfm4* expression levels after 6Gy, (**h**) *Olfm4* expression levels after 8Gy, (**i**) *Alpi1* expression levels after 6Gy, (**j**) *Alpi1* expression levels after 8Gy, (**k**) *Chga* expression levels after 6Gy, and (**l**) *Chga* expression levels after 8Gy were calculated as a fold change at 6, 24, 48 and 72 h with either ^137^Cs or X-ray irradiation. *Actb* was used as a housekeeping gene (control). Data points represent the average of three independent experiments, with the mean ±SD indicated. Significance was determined by the student’s test followed by an analysis of the normal distribution (Tukey’s test), *p < 0.05, **p < 0.01, ***p < 0.001, ****p < 0.0001.

### Mouse models

*Bmi1-Cre^ER^* mice were exposed to the irradiation as described in the Materials and Methods section and as depicted in Supplementary Figure 1c. In addition to assessing potential differences between 12Gy total body irradiation (TBI) using ^137^Cs and X-ray irradiation, we include 12Gy abdominal (ABD) irradiation with X-ray irradiation to localize damage toward a specific organ, such as intestine. Representative H&E images of duodenum, jejunum, and ileum of *Bmi1-CreER* are shown in Figure 5. ^137^Cs TBI, X-ray TBI, and X-ray ABD irradiation resulted in shortening of the villi and loss of crypts at 24-48h and enlarged proliferated crypts at 96h post irradiation (Figure 5). One key feature of intestinal regeneration after injury is the enlargement of the crypts with still shortened villi, which are very well defined at 96 hours after irradiation^32^. We observed that at 96h, the villi are significantly shortened under all three irradiation conditions, with abdominal irradiation demonstrating a slight increase in villi length compared to 12Gy TBI, irrespective of the source (Supplementary Figure 3a). In contrast, under all three tested conditions, the crypts’ length showed significant enlargement, without significant differences based on source or type of irradiation (Supplementary Figure 3b). Similarly, in the villi-shortening condition, the crypt/length axis was significantly decreased in three tested conditions, with the villi/crypt’s axis being significantly longer upon ABD irradiation compared to TBI, regardless of the source of irradiation (^137^Cs or X-ray) (Supplementary Figure 3c).

**Figure 5.**
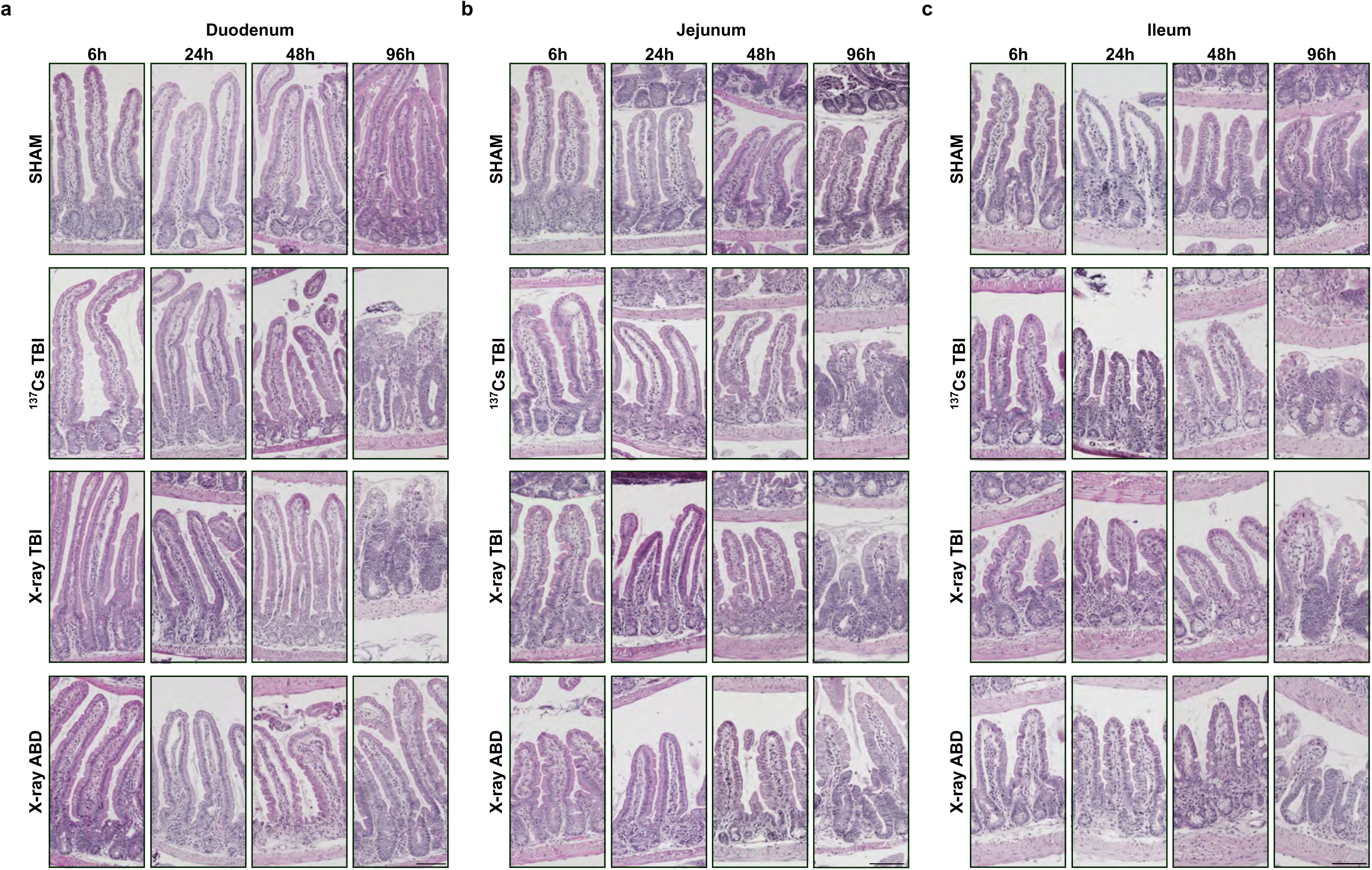
Intestinal epithelium regeneration in *Bmi1-Cre^ER^* mouse after 12Gy TBI with ^137^Cs, X-ray, or 12Gy X-ray ABD irradiation. Representative images of hematoxylin and eosin staining in the duodenum (**a**), jejunum (**b**), and ileum (**c**) of *Bmi1-Cre^ER^* mice exposed to 0 or 12 Gy total body irradiation (TBI) at 6-, 24-, 48-, and 96-h using ^137^Cs, X-ray, or X-ray ABD irradiation. Scale bar = 100 µm.

Previous studies from our lab and others showed the almost complete absence of p21 in intestinal crypts during homeostasis, with sharply induced levels upon injury using ^137^Cs irradiator^33–35^. Similarly, the intestinal crypts are characterized by low levels of p53 at the steady state; however, upon radiation injury, p53 levels increase specifically at early time points (24-48h)^33,36,37^. We and others also showed that MSI1 expression is low in the intestinal epithelium during homeostasis and increases upon ^137^Cs TBI irradiation^38–40^. Thus, we performed immunofluorescence (IF) staining and analyzed the expression patterns of p21, MSI1, and p53 in the duodenum at 96 hours post-irradiation. We observed a similar pattern of p21 and MSI1 expression post-irradiation, regardless of the irradiation source (Figure 6a-b). However, we observed lower MSI1 expression in the duodenum at 96h post-X-ray ABD compared to both TBI conditions (Figure 6c). The p53 expression pattern at 24h and 48h post-irradiation with ^137^Cs and x-ray irradiators was similar (Supplementary Figure 4a and b). However, we observed lower p53 expression at 48h after X-ray ABD (Supplementary Figure 4c), suggesting that the tissue recovers faster with X-ray ABD than with other treatments.

**Figure 6.**
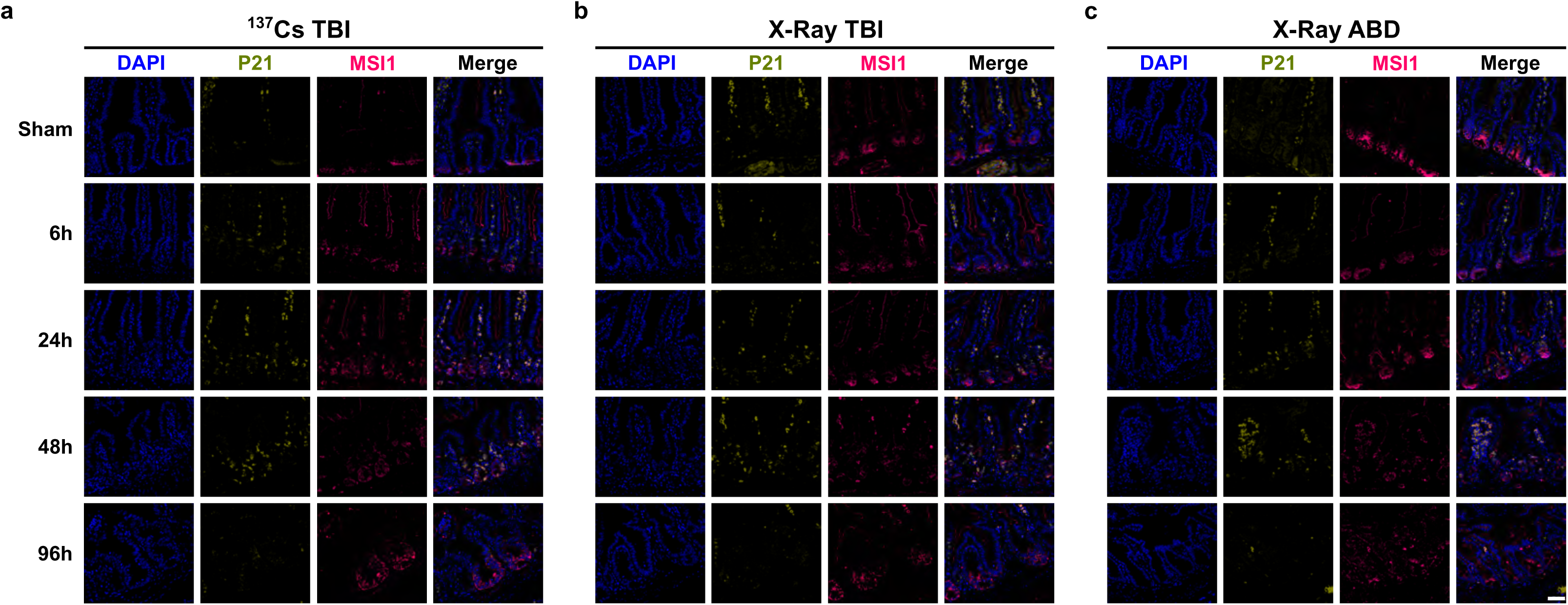
Time-dependent P21 and MSI1 expression pattern in the intestinal epithelium of *Bmi1-Cre^ER^*mouse after 12Gy TBI with ^137^Cs, X-ray, or 12Gy X-ray ABD irradiation. Representative images of immunofluorescence staining for P21, MUSHASHI-1 (MSI1), and nuclei marker (DAPI) in the duodenum of *Bmi1-Cre^ER^*mice exposed to 0 or 12 Gy TBI at 6-, 24-, 48-, and 96-h using ^137^Cs (**a**), or X-ray irradiator (**b**), or 12Gy X-ray ABD irradiation (**c**). Scale bar = 100 µm.

We also performed immunohistochemical (IHC) staining for OLFM4 and SOX9 in the duodenum of collected tissues at 6, 24, 48, and 96h post-irradiation. OLFM4, a stem cell marker^41^, is highly expressed at the base of the crypts in non-irradiated mice. In contrast, it gradually decreases over time, with complete abolition at 48h post-irradiation, regardless of the type of irradiation (Figure 7a). Compared to the sham, the number of OLFM4-positive crypts was diminished 96h post-irradiation, with the abdominal X-ray injury surpassing levels under whole-body irradiation regardless of the injury condition (Figure 7a and Supplementary Figure 3d). SOX9 is an intestinal crypt transcription factor, target of WNT signaling, and its level increased during the regenerative phase post-injury^18,42^. We observed a gradual decrease in SOX9 expression at 6, 24, and 48 h post-irradiation. While there are almost no differences in the number of SOX9-positive cells per regenerative crypt between TBI with ^137^Cs and X-ray, there are fewer of them than in the ABD irradiation (Figure 7b and Supplementary Figure 3e). Our laboratory has shown that the reserved stem cell subpopulation, derived from BMI1-positive cells, plays an important role in regeneration, as it actively proliferates and replenishes intestinal epithelial crypts, allowing the gut to return to homeostasis^19,38,43^. To assess the impact of different irradiation conditions on the regenerative capacity of BMI1-positive cells, we irradiated tamoxifen-treated *Bmi1-Cre^ER^*mice. We counted BMI1(YFP)-positive and MKi67-positive crypts in the duodenum at 96h post-irradiation (Figure 8a). We observed an increase in the average number of YFP-positive crypts at 96h with X-ray abdominal irradiation compared to other irradiation conditions (^137^Cs TBI and X-ray TBI) (Figure 8b). However, we did not observe a significant difference in the percentage of MKi67-positive cells between sham and irradiated tissues at 96h post-irradiation (Figure 8c).

**Figure 7.**
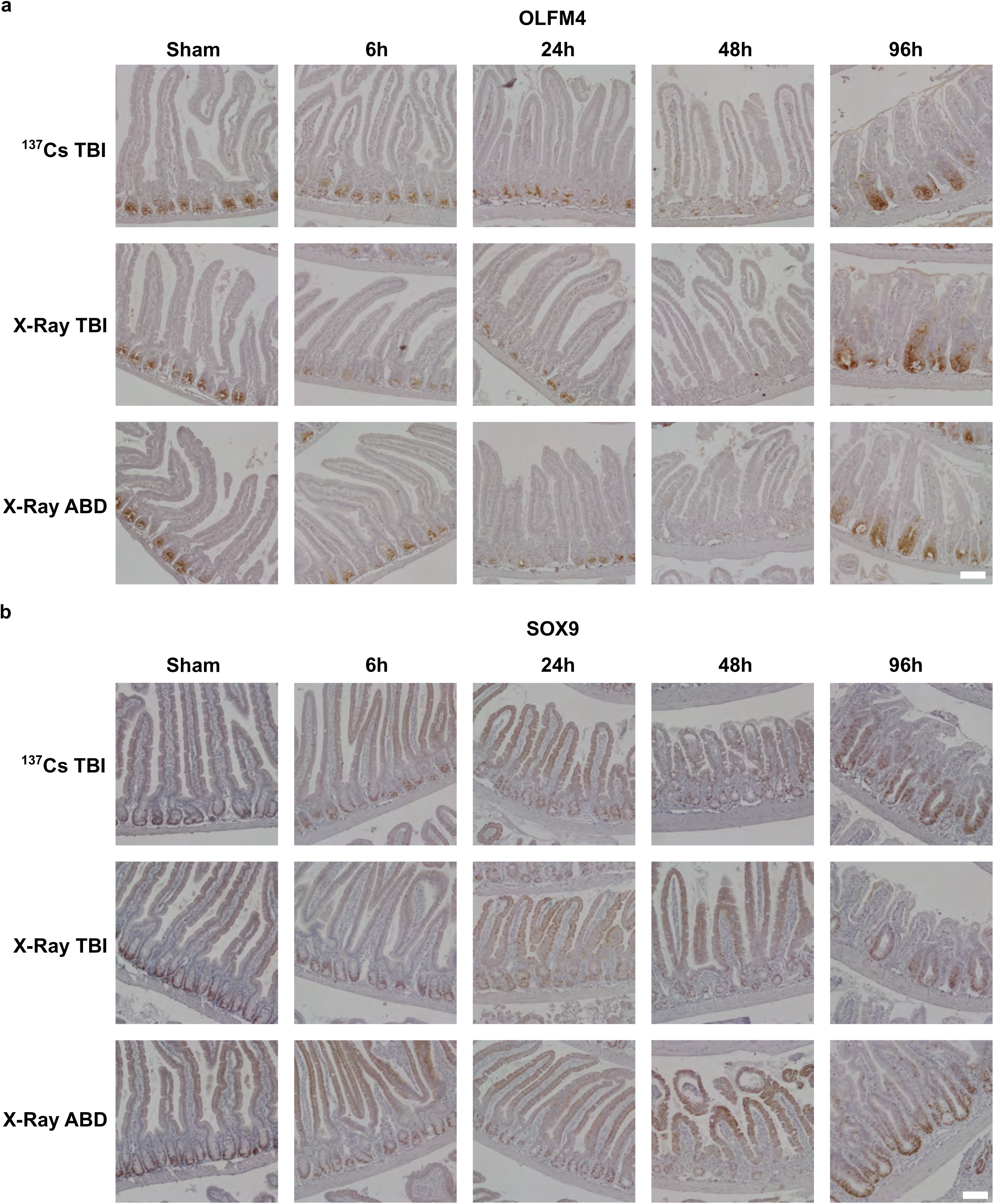
Time-dependent OLFM4 and SOX9 expression pattern in the intestinal epithelium of *Bmi1-Cre^ER^*mouse after 12Gy TBI with ^137^Cs, X-ray, or 12Gy X-ray ABD irradiation. Representative images of immunohistochemical staining for OLFM4 (**a**) and SOX9 (**b**) in the duodenum of *Bmi1-Cre^ER^*mice exposed to 0 or 12 Gy TBI at 6-, 24-, 48-, and 96-h using ^137^Cs irradiator, or X-ray irradiator, or 12Gy X-ray ABD irradiation. Scale bar = 100 µm.

**Figure 8.**
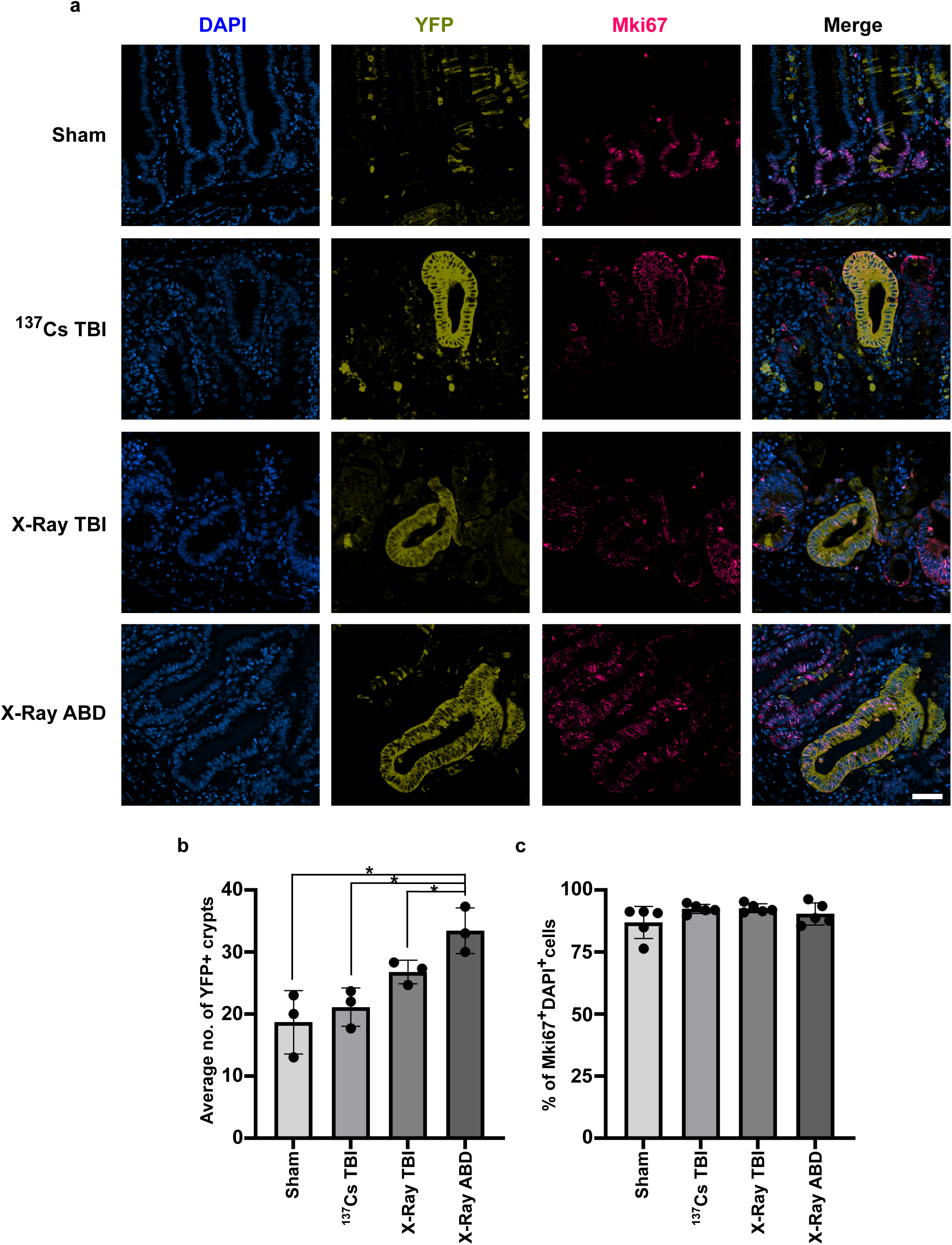
YFP and MKi67 expression pattern in the intestinal epithelium of *Bmi1-Cre^ER^* mouse after 12Gy TBI with ^137^Cs, X-ray, or X-ray ABD irradiation. Representative images of immunofluorescence staining (**a**) for *Bmi1^YFP^*-positive cells marker (YFP), proliferation marker (MKi67), and nuclei marker (DAPI) in the duodenum of *Bmi1-Cre^ER^* mice exposed to 0 or 12 Gy TBI at 96 h with ^137^Cs or X-ray irradiation or 12Gy X-ray ABD irradiation. Scale bar = 50 µm. Quantification of proliferating crypts (**b**) of the duodenum from mice exposed to 0 or 12 Gy ^137^Cs or X-ray TBI or 12Gy X-ray ABD irradiation at 96 h. Analysis was performed on sham (100 crypts) and irradiated (300 crypts) mice. Quantification of MKi67-positive and DAPI-positive crypts of the duodenum from mice exposed to 0 or 12 Gy ^137^Cs or X-ray TBI or 12Gy X-ray ABD irradiation at 96 h. Data points represent the average of three (**b**) or five (**c**) independent experiments with the mean ±SD indicated. Significance was determined by the Student’s test followed by an analysis of the normal distribution (Tukey’s test), *p < 0.05, **p < 0.01, ***p < 0.001, ****p < 0.0001.

## Discussion

Radiation injury in mice and rats is a useful tool for studying the intestinal response to injury and the gut’s ability to regenerate. The majority of studies to date use 10-15Gy whole body irradiation with ^137^Cs as a source of injury. This model is characterized by early injury within 24-48 hours, followed by a proliferative/regenerative phase leading to recovery, and allows the establishment of mechanisms governing intestinal regeneration. Recent studies extend the use of radiation to induce injury in organoids as an alternative to animal models. Furthermore, decades of ^137^Cs irradiation have been used to study the mechanisms governing colorectal cancer cell lines in standard 2D models. Due to the recent transition from ^137^Cs to X-ray irradiators, several laboratories have begun analyzing potential differences between these two sources of irradiation; most focusing on the assessment of hematology and blood chemistry. Our laboratory has used ^137^Cs irradiation for decades to induce injury in colorectal cancer cell lines and mouse models, and we decided to assess whether there are macroscopic biological differences between the two sources of delivered radiation, as our institution was planning to decommission the ^137^Cs irradiator in Spring 2025. In this study, we evaluated the effects of both irradiation sources on colorectal cancer cell lines, mouse intestinal organoids, and mouse intestinal epithelium.

Consistent with our hypothesis, we have not observed any significant difference in the response of the three tested colorectal cancer cell lines regardless of the source of irradiation. We assessed markers of injury and proliferation, both on RNA and protein levels, and we obtained very similar results. In contrast, while working with organoids, we obtained results that differed for some of the tested factors, specifically those connected with cell proliferation, stem cell function, and differentiation. We observed more robust increases in the RNA levels of these factors (*Mki67*, *Lgr5*, and *Olfm4*) at the later time points after irradiation (48 and 96h) whether we used 6 or 8Gy dose. This observation suggests that the organoids under tested conditions recover faster after X-ray irradiation compared to ^137^Cs. Similarly, we have noticed slight variations in the levels of two tested lineage markers – *Alpi* and *Chga* between both tested sources of irradiation. These results warrant the need of more comprehensive studies and diligent analysis while comparing studies using various irradiation sources.

Further, we assessed the impact of two different sources of irradiation in mice. For this set of experiments, we added abdominal irradiation combined with X-ray irradiation, as this equipment allowed for more precise irradiation. All three experimental settings caused injury to the intestinal epithelium that was followed by regeneration. Consistent with the expectations, a difference occurred between TBI and ABD irradiation models, with the latter showing less shortening of villi and crypt/villi axis and larger number of YFP-positive crypts or fewer SOX9-positive cells in the crypts 96h post-treatment.

In summary, we compared the effects of irradiation delivered by ^137^Cs and X-ray sources in three biological models and obtained comparable results with both sources, with slight differences in the levels of specific markers in all tested models. Besides reasonably similar results from both irradiators, there are clear advantages associated with X-ray irradiation. X-ray irradiators permit extremely accurate delivery of radiation not only to the whole animal but also to specific organs, increasing the precision of the irradiation and leading to greater reproducibility of experiments. The experiments performed here have several limitations. Within the allotted time before the equipment’s decommissioning, we tested only three colorectal cancer cell lines, mouse organoids from a single source, and two mouse strains. The performed analysis was limited and focused on a subset of targets, as we could not test all genes, proteins, and changes across all pathways and cell populations *in vitro* and *in vivo*.

## Methods

### Cell lines

Human colorectal cancer (CRC) cell lines DLD-1 (ATCC, CCL-221, RRID: CVCL_0248), HCT116 (ATCC, CCL-247, RRID: CVCL_0291), and RKO (ATCC, CRL-2577, RRID: CVCL_0504) were obtained from the American Type Culture Collection (ATCC; Manassas, VA). DLD-1 cells were cultured in RPMI-1640 medium (Corning, 10–040-CV); RKO cells in DMEM medium (Corning, 10–013-CV); and HCT116 cells in McCoy’s 5a medium (Corning, 10–050-CV). All media were supplemented with 10% fetal bovine serum (Peak Serum, PS-FB3) and 1% antibiotic/antimycotic (10,000 μg/mL penicillin and 10,000μg/mL streptomycin; Gibco, 15140–122). Cell lines were incubated and maintained at 37°C with 5% CO_2_, in a humidified atmosphere. *In vitro* experiments were conducted at cell passages lower than 30^44^.

### Mice

*Bmi1-Cre^ER^*;*Rosa26^eYFP^* (*Bmi1-Cre^ER^*) mice were described previously^38,45,46^. The mice were given standard chow and water ad libitum. Transnetyx performed the genotyping. Both male and female mice, 8 to 12 weeks old, were used in this study. Animal studies are approved by the Stony Brook University Institutional Animal Care and Use Committee under protocol IACUC2023-00041.

### Crypt isolation for enteroids culture

The mouse intestinal crypts were isolated using previously established protocol^47^ and grown in Matrigel droplets and maintained in L-WRN conditioned media. Organoid culture media was obtained from L-WRN cell line culture as previously described^48^ and supplemented with 1 x N2 supplement (Thermo Fisher Scientific, 17502-048), 1 x B27 supplement (Thermo Fisher Scientific, 17504-044), 10nM gastrin I (Sigma-Aldrich, G9145), 50ng/mL recombinant human Epidermal Growth Factor (Thermo Fisher Scientific, C-60169A), 500nM transforming growth factor β inhibitor A83-01 (Tocris Bioscience, 2939), 1mM N-acetylcysteine (Sigma-Aldrich, A9165), 100µg/mL Primocin (Invitrogen, ant-pm-2). During the first two days of culture, the media was also supplemented with 10µM GSK3β inhibitor CHIR99021 (Tocris, 4423) and 10µM ROCK inhibitor Y-27632 (Sigma-Aldrich, Y05030).

### Radiation experiments

^137^Cs irradiation of cell lines, organoids, and 8-12-week-old mice was administered at a dose rate of 0.759 Gy/min. The Xstrahl XenX (serial number X2023-1013) was used for X-ray irradiation of cell lines, organoids and 8-12-week-old mice using 220kV X-rays and 0.15mm copper. A dose of 12Gy to cell lines and 6 or 8Gy to organoids were delivered at a dose-rate of 3.18Gy/min. Additionally, a dose of 12Gy was delivered to mice either at a dose-rate of 3.18Gy/min for abdominal irradiation or at a dose-rate of 0.5811Gy/min for total body irradiation.

CRC cell lines were seeded in 6-well plates at a concentration of 5 x 10^5^ cells/well in 2mL of the appropriate media. Twenty-four hours later, cells were irradiated with 12Gy using either a ^137^Cs-or an X-ray irradiator (XenX Xstrahl). The cells were collected at 6, 24, 48, and 72 h and used for RNA and protein extraction (Supplementary Figure 1a). RT-qPCR was used to analyze gene expression, and Western blot was used to analyze protein levels. Each experiment was performed in triplicate.

Mouse intestinal enteroids were seeded in 24-well plates in Matrigel and 500μL of L-WRN media described above. Twenty-four hours later, enteroids were irradiated with 6Gy or 8Gy either with a ^137^Cs irradiator or an X-ray irradiator (XenX Xstrahl). Enteroids were imaged at 6-, 24-, 48-, and 72-hours post-radiation and collected for total RNA and protein extraction (Supplementary Figure 1b). RT-qPCR was used to analyze gene expression, and Western blot was used to analyze protein levels. Each experiment was performed in triplicate.

For mouse experiments, tamoxifen (Sigma-Aldrich, T5648) was dissolved in corn oil (30 mg/mL) and administered as a single intraperitoneal injection (225 mg/kg) 2 days before exposure to radiation. The experimental groups of *Bmi1-Cre^ER^* and *Pdgfr-Cre;mTmG* mice were exposed either to 12 Gy γ irradiation from ^137^Cs at a dose rate of 0.759 Gy/min, X-ray TBI, or to X-ray abdominal irradiation (ABD). The control (sham) groups received 0Gy radiation. The mice were euthanized, and the small intestine was collected at 6, 24, 48, and 96 h after irradiation (Supplementary Figure 1c). All mice were injected intraperitoneally with 100μg of EdU (Santa Cruz Biotechnology, sc-284628) dissolved in 1:5 (v/v) DMSO and sterile water 3 h before collection. Proximal, middle, and distal parts of the small intestine were collected using the previously described Swiss-rolled technique ^49,50^.

### Hematoxylin and Eosin staining

Formalin-fixed, paraffin-embedded (FFPE) blocks were sectioned at 5μm, dried, and baked overnight at 65°C in an oven. The next day, the slides were cooled for 10 min, deparaffinized in 100% xylene for 3 min (2 changes), and rehydrated in a decreasing ethanol gradient (100% ethanol, 2 min; 95% ethanol, 2 min; 70% ethanol, 2 min). Slides were then rinsed in running distilled water for 2 min, stained with Gill’s III hematoxylin for 2 min, and rinsed in running tap water for 5 min. Next, the slides were dipped 10 times into a 5% (w/v) lithium carbonate solution and rinsed in running distilled water for 2 min. Then, the slides were stained with eosin Y with phloxine for 5 min and briefly rinsed in 70% ethanol. Finally, slides were dehydrated in an increasing ethanol gradient: 95% ethanol, 5 dips, and 100% ethanol, 5 dips, and cleared in two changes of 100% xylene for 2 min. The slides were mounted using Cytoseal XYL Mounting Medium (Thermo Fisher Scientific, 8312-4). The images were obtained using an Eclipse 90i microscope (Nikon, Tokyo, Japan) equipped with a DS-Fi1 camera (Nikon).

### Western Blot

Protein samples were lysed in 2× Laemmli buffer. Proteins were electrophoresed on 4%–20% polyacrylamide gels (Bio-Rad, 5671094) and transferred to nitrocellulose membranes (Bio-Rad, 162-0097). The membranes were blocked with 5% nonfat milk in 1× Tris-buffered saline containing 0.01% Tween-20 (TBST) for 1 hour at room temperature and incubated on a rocking platform overnight with primary antibodies against phospho-β-catenin Ser552 (1:1000; Cell Signaling Technology, 9566S), β-catenin (1:1000; Cell Signaling Technology, 8480P), phospho-TP53 Ser15 (1:1000; Cell Signaling Technology, 9284), TP53 (1:1000; Cell Signaling Technology, 9282), KLF5 (1:1000; Abcam, 137676), BAX (1:1000; Cell Signaling Technology, 89477), γ-H2AX (1:2000; Cell Signaling Technology, 9718), p21 (1:1000; BD Biosciences, 556430), and β-actin (1:2000; Sigma-Aldrich, A1948). The membranes were then washed in 1× TBST, incubated with appropriate IgG secondary antibodies conjugated with horseradish peroxidase for 1 hour at room temperature, and developed using SuperSignal West Pico PLUS Chemiluminescent Substrate (Thermo Fisher Scientific, 34080) and Immobilon Western Chemiluminescent HRP Substrate (Millipore Sigma, WBKLS0500) using Azure400 Gel Imaging System (Azure Biosystems, Dublin, CA). The membranes were probed for β-actin as an internal control. Loading controls were run on the same blots. The experiments were performed in 3–5 independent biological replicates, as indicated in the figure legends.

### Immunofluorescent staining (IF)

Proximal parts of the small intestine were collected using the Swiss-rolled technique as described previously^43,49^. Paraffin-embedded blocks were cut into 5μm thick sections, dried, and baked overnight at 65°C in an oven. The next day, slides were cooled for 10 min, deparaffinized in 100% xylene for 3 min (2 changes), incubated at room temperature in 2% hydrogen peroxide in methanol for 30 min, and rehydrated in ethanol gradient (100% ethanol, 2 min; 95% ethanol, 2 min; 70% ethanol, 2 min.). Antigens were retrieved by incubating the slides in 10mM Na-citrate buffer (pH 6.0) at 110°C for 20 min in a pressure cooker. The sections were then washed with distilled water, incubated for 1 h at 37°C in a blocking solution (5% BSA in TBST), and incubated with primary antibodies against GFP (1:500, AvesLabs, GFP-1010), MSI1 (1:200, MBL International Corporation, D270-3), MKi67 (1:300, DAKO, M7249), P21 (1:200, BD Biosciences, 556430), and P53 (1:200, Invitrogen, MA5-50452) on a rocking platform overnight at 4°C. The next day, slides were washed three times with TBST and incubated with appropriate secondary antibodies conjugated with fluorophores for 30 min at 37° C in a blocking solution. The tissues were also counterstained with Hoechst 33258 (1:1000, Invitrogen, H3569) to visualize the nuclei. Slides were mounted with Fluorescent Mounting Media, Aqueous (Millipore, Sigma). The images were obtained using an Eclipse 90i fluorescence microscope (Nikon) equipped with a DS-Qi1Mc camera (Nikon).

### Immunohistochemical Staining (IHC)

On the first day of Immunohistochemical staining (IHC), the same steps were performed as for IF for both OLFM4 and SOX9, except that for SOX9 staining, the slides were incubated in 2% hydrogen peroxide in methanol for 30 min at room temperature after antigen retrieval. The sections were then incubated with primary antibodies against OLFM4 (1:1000 Cell Signaling, 39141T) and SOX9 (1:3000 Sigma, AB5535) overnight at 4°C. The next day, slides were washed three times with TBST and OLFM4 slides were incubated with secondary antibody-HRP probe (Biocare Mach 3 Rabbit HRP polymer kit) for 30 min at 37°C followed by tertiary antibody-HRP probe (Biocare Mach 3 Rabbit HRP polymer kit) for 30 min at 37°C. For SOX9, slides were incubated with appropriate horseradish peroxidase-conjugated secondary antibodies for 1 h at 37°C, then washed with TBST. Both OLFM4 and SOX9 slides were then washed with TBST and incubated with DAB reagent (Betazoid Dab chromogen kit, BDB2004L), followed by another wash in distilled water. Next, slides were counterstained with hematoxylin solution for 1 min and rinsed in running tap water for 2 min. Next, the slides were dipped 10 times into a 5% (w/v) lithium carbonate solution and rinsed in running distilled water for 2 min. Finally, slides were dehydrated in an increasing ethanol gradient: 95% ethanol, 5 dips, and 100% ethanol, 5 dips, followed by clearing in 2 changes of 100% xylene for 3 min. The slides were then mounted using Cytoseal XYL Mounting Medium (Thermo Fisher Scientific, 8312-4). The images were obtained using an Eclipse 90i microscope (Nikon, Tokyo, Japan) equipped with a DS-Fi1 camera (Nikon).

### RNA isolation and Gene Expression Analysis of Cells by RT-qPCR

RNA was isolated using the RNeasy Mini Kit (Qiagen, 74106). RNase-Free DNase Set (Qiagen, 79254) was used to remove genomic DNA. The RNA was examined for purity and concentration using a NanoDrop spectrophotometer (NanoVue Plus; GE Healthcare, Milwaukee, WI). First-strand complementary DNA (cDNA) was synthesized using 1μg of total RNA and the SuperScript VILO cDNA Synthesis Kit (Thermo Fisher Scientific, 11756050). The reaction was performed according to a standard protocol: 10 min at 25°C, followed by 10 min at 50°C, and 5 min at 85°C in a Mastercycler X50s system (Eppendorf, Hamburg, Germany). RT-qPCR analysis was performed using TaqMan Gene Expression Master Mix (Thermo Fisher Scientific, 43-690-16) and QuantStudio 3 (Applied Biosystems, Waltham, MA) with 10 min at 95°C followed by 40 cycles of 15 sec at 95°C and 1 min at 60°C. Commercially available TaqMan primers detecting mouse *ChgA* (Mm00514341-FAM), *Alpi1* (Mm01285814-FAM), *Olfm4* (Mm01320260-FAM), *Cdkn1a* (Mm00432448-FAM), *Lgr5* (Mm00438890-FAM), *Mki67* (Mm01278617-FAM), *Actb* (Mm04394036-VIC) and human *CDKN1A* (Hs00355782-FAM), *KLF4* (Hs00358836-FAM), *KLF5* (Hs0000000-FAM), *BCL2* (Hs01048932-FAM), *BAX* (Hs00180269-FAM) and *HPRT1* (Hs02800695-VIC) transcripts were used. All kits were used according to the manufacturer’s instructions. The experiments were performed in 3–5 independent biological replicates.

### Crypt/villi axis scoring

For crypt regeneration scoring, fragments corresponding to 20 crypts were quantified from each specimen collected from 12Gy ^137^Cs or X-ray TBI, 12Gy ABD, or sham irradiated mice. The lengths of the crypts and villi were measured in twenty consecutive crypts at 96 h post-irradiation.

### YFP^+^, OLFM4^+^, SOX9^+^, MKi67^+^, and DAPI counts

For YFP-positive and OLFM4-positive counts, 100 crypts per sham-irradiated and 300 crypts per 12Gy ^137^Cs or X-ray TBI, or 12Gy X-ray ABD mouse were quantified at 96 h. For MKi67- and DAPI-positive cell counts and SOX9-positive cell counts, 20 crypts were quantified per mouse.

### Statistical Analysis

The analysis was performed using an appropriate statistical test with a value of p < 0.05 considered significant using GraphPad Prism version 10 for Windows (GraphPad Software, Inc., San Diego, CA).

## Acknowledgements

The authors acknowledge the Stony Brook Cancer Center Research Histology Core for assistance with biospecimen preparation. This research was funded by the National Institutes of Health grant DK052230 awarded to VWY, and this material is based on work supported by the U.S. Department of Energy, National Nuclear Security Administration, under Contract DE-AC05-76RL01830, with support awarded to ABB.

## Author contributions

Conception: VWY and ABB; design: ABB; acquisition: RL, EJO, SK, ZJW, MR, and MG; analysis: RL, EJO, SK, ZJW, MR, and MG; interpretation of data: RL, EJO, SK, ZJW, MR, MG, VWY, and ABB; writing – original draft, review, and editing: RL, EJO, SK, ZJW, MR, MG, VWY, and ABB.

## Competing interests

The author(s) declare no competing interests.

## Data availability statement

All relevant data is contained within the article. The original contributions presented in the study are included in the article/supplementary material; further inquiries can be directed to the corresponding author.

## Supplementary figure legends

**Supplementary Figure 1. In vitro and in vivo experimental outline.** (**a**) – colorectal cancer cells (HCT116, RKO, and DLD-1) were seeded in a 6-well plate, and 24 h later, exposed to 12Gy irradiation with ^137^Cs or X-ray irradiator. Samples for analysis were collected at 0, 6, 24, 48, and 72 h post-irradiation. (**b**) – Mouse intestinal enteroids were seeded in a 24-well plate, and 24 h later, exposed to 6Gy or 8Gy irradiation with ^137^Cs or X-ray irradiator. Samples for analysis were collected at 0, 6, 24, 48, and 96 h post-irradiation. (**c**) – Eight-twelve week old mice were exposed to 12Gy TBI with ^137^Cs or X-ray irradiator, or 12Gy ABD with X-ray irradiator. Samples for analysis were collected at 0, 6, 24, 48, and 96 h post-irradiation (created with Biorender).

**Supplementary Figure 2. Quantification of western blot analysis of markers of cell injury, proliferation, and survival in three colorectal cancer cell lines presented in Figure 2**. Markers of cell injury (p21, γH2AX, TP53, and phospho-TP53), proliferation (KLF5), and pro-apoptotic (BAX) markers were analyzed using Western blot and densitometry analyses was performed with ImageJ software. HCT116 (**a**), RKO (**b**), and DLD-1(**c**). Data points represent the average of three independent experiments, with the mean ±SD indicated. Significance was determined by the student’s test followed by an analysis of the normal distribution (Tukey’s test), *p < 0.05, **p < 0.01, ***p < 0.001.

**Supplementary Figure 3. Assessment of duodenum at 96h post irradiation using 12Gy TBI ^137^Cs or X-ray irradiation, or 12Gy X-ray ABD irradiation.** Analysis of villi length (**a**), crypt length (**b**), villi/crypt axis length (**c**), OLFM4-positive (**d**) and SOX9-positive (**e**) crypts in 12Gy X-ray ABD irradiated tissues compared to ^137^Cs and X-ray irradiated tissues. Data points represent the average of five independent experiments, with the mean ±SD indicated. Significance was determined by the student’s test followed by an analysis of the normal distribution (Tukey’s test), *p < 0.05, **p < 0.01, ***p < 0.001, ****p < 0.0001.

**Supplementary Figure 4. Time-dependent p53 expression pattern in the intestinal epithelium of *Bmi1-Cre^ER^* mouse after 12Gy TBI with ^137^Cs or X-ray irradiation or X-ray ABD irradiation**. Representative images of immunofluorescence staining for p53 and nuclei marker (DAPI) in the duodenum of *Bmi1-Cre^ER^* mice exposed to 0 or 12 Gy TBI at 6-, 24-, 48-, and 96-h using ^137^Cs, or X-ray irradiator or 12Gy X-ray ABD irradiation. Scale bar = 100 µm.

## Notes

### Competing Interest Statement

The authors have declared no competing interest.

