## Supplementary figures and images for "Comparison studies between Cesium-137 and X-ray irradiators in epithelial injury using *in vitro* and *in vivo* models"

### Supplementary Figure 1

a

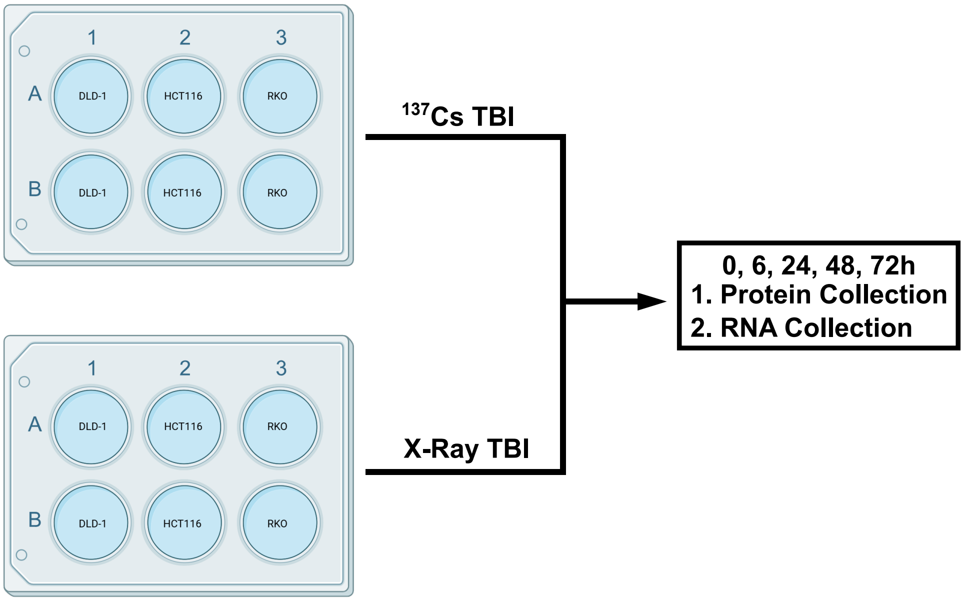


b

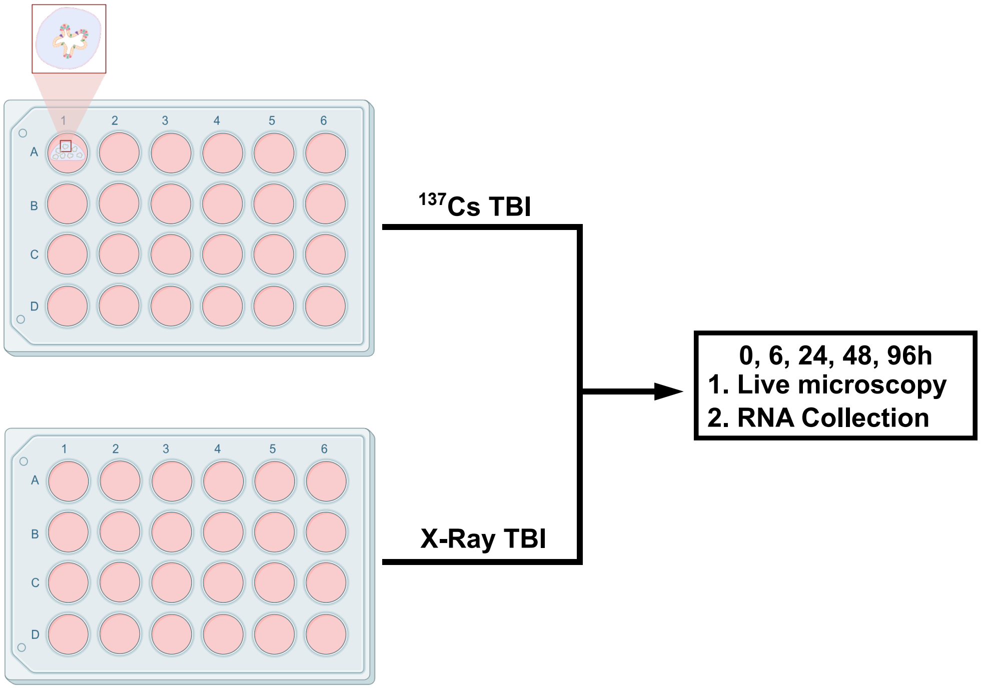


c

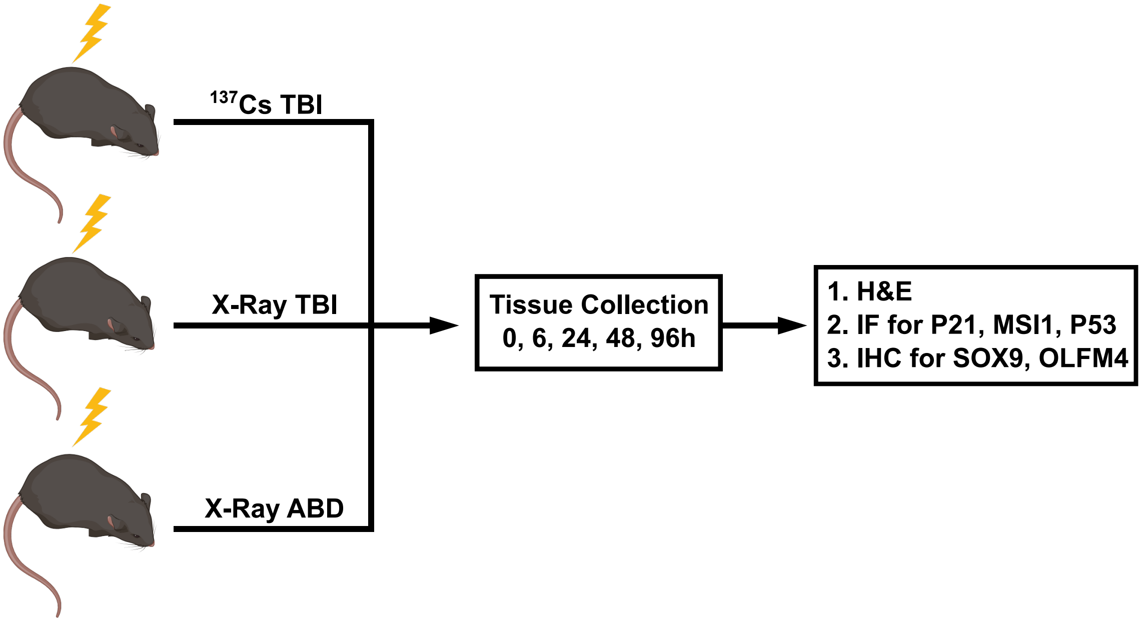


**Supplementary Figure 1.**

### Supplementary Figure 2

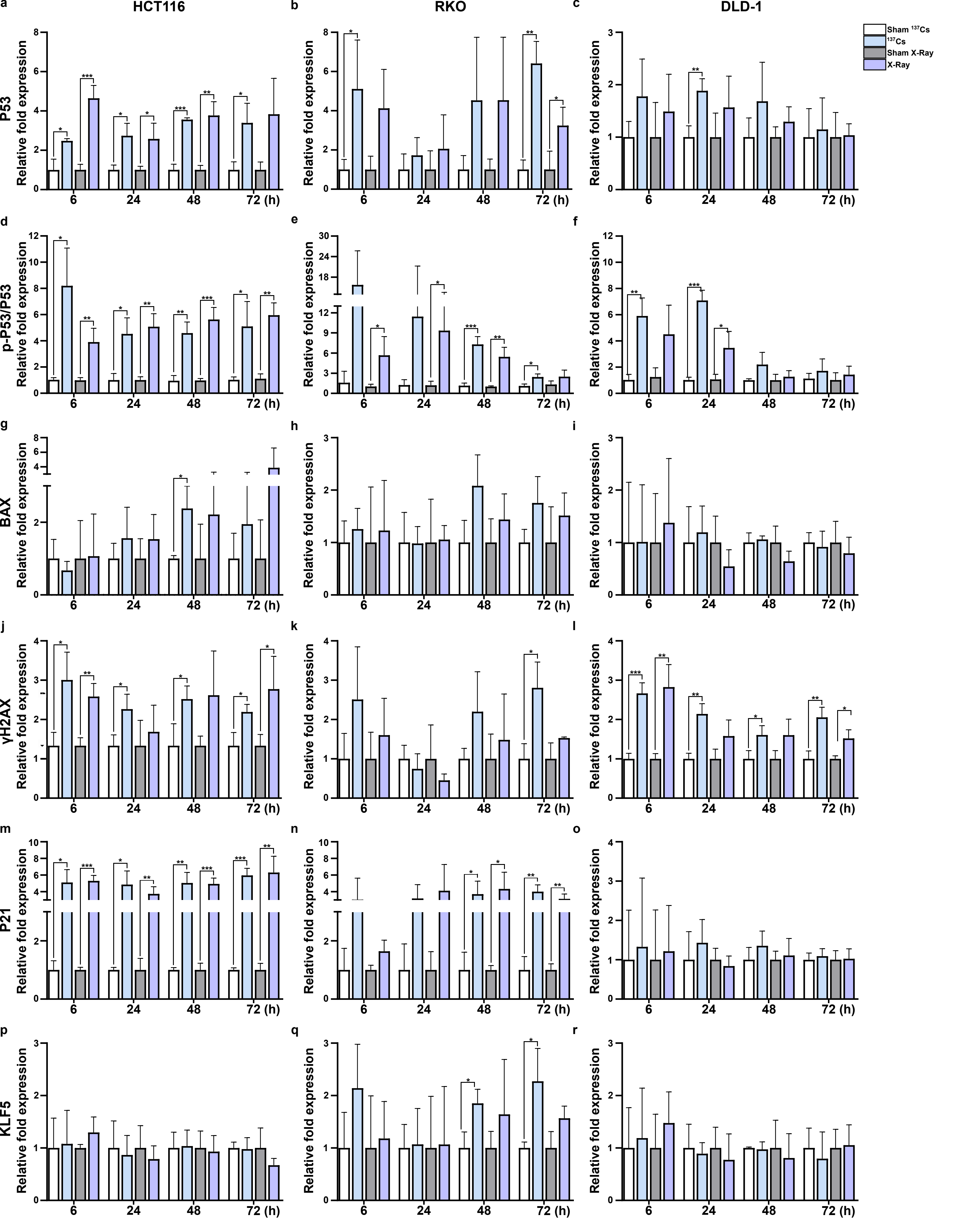


Supplementary Figure 2.

### Supplementary Figure 3

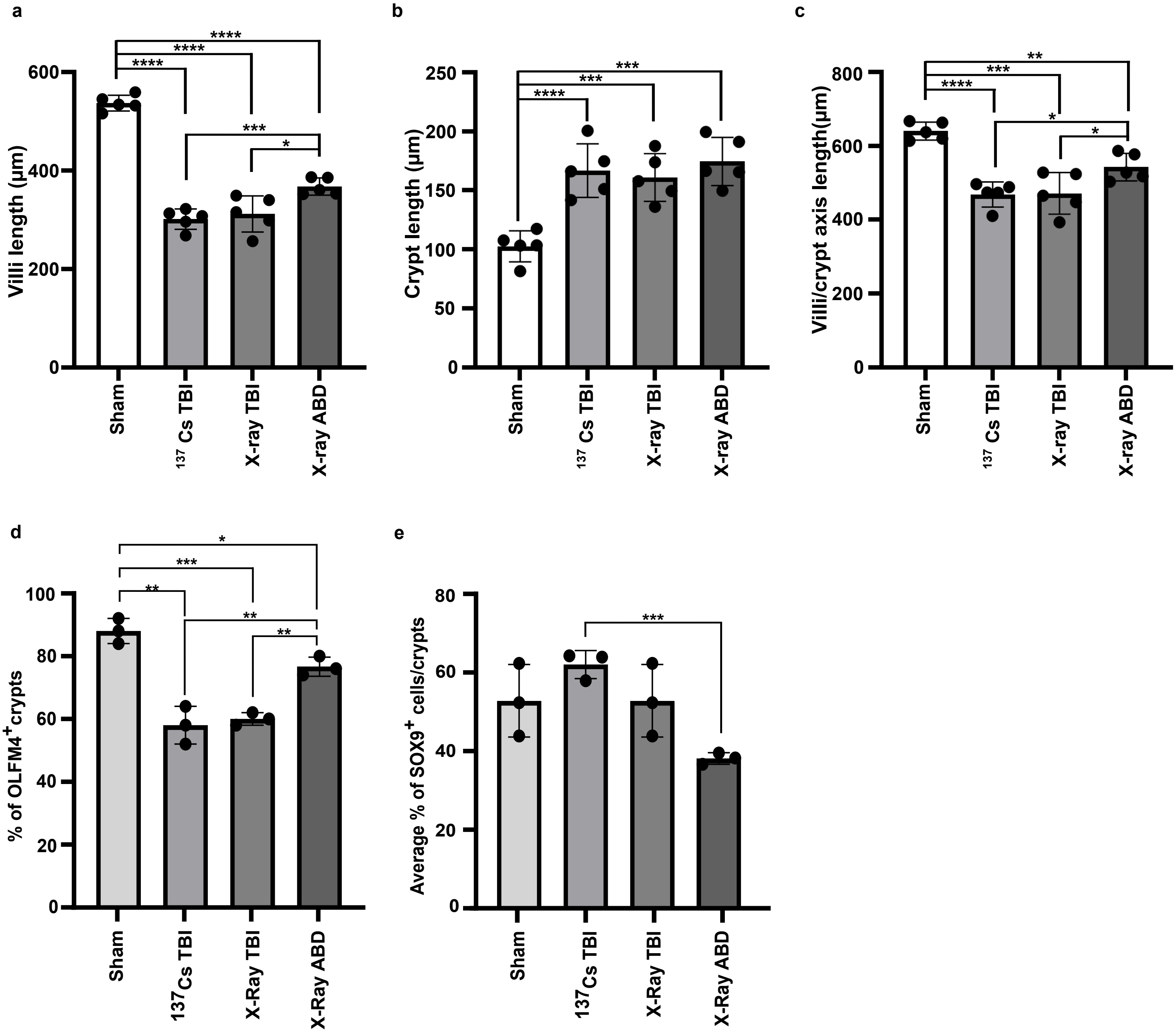


**Supplementary Figure 3.**

### Supplementary Figure 4

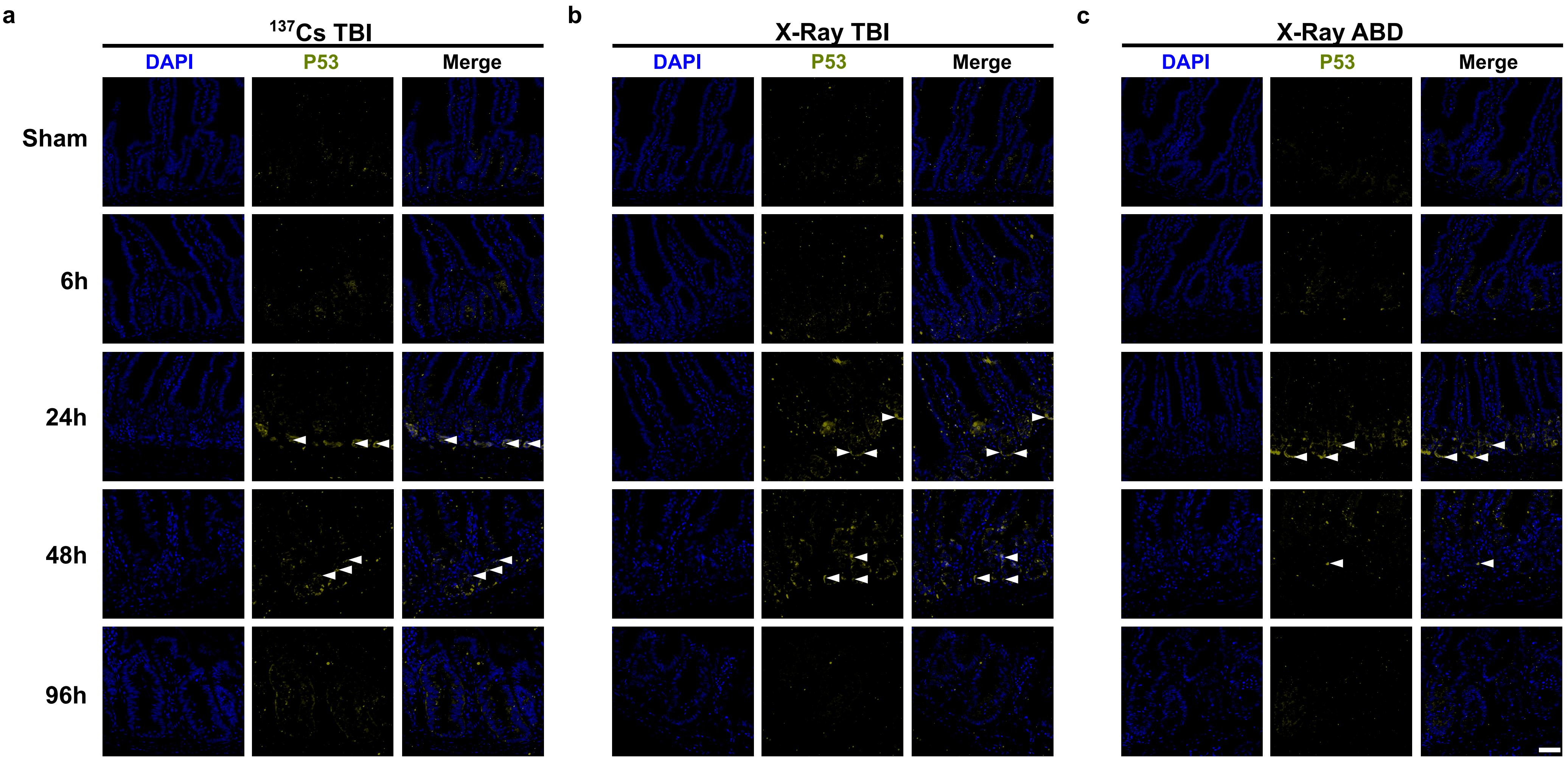


**Supplementary Figure 4.**
